# Multispecies transport shapes electrochemical environments in perforated cells

**DOI:** 10.64898/2026.11.03.764109

**Authors:** Yikang Guan, Hengyu Li, Miao Shi, Kenneth Sajbel, Yi Byron Gao, Shiyu Xia

**Affiliations:** Department of Chemical and Biomolecular Engineering, College of Chemistry, University of California, Berkeley, Berkeley, CA 94720, USA; Division of Chemistry and Chemical Engineering, California Institute of Technology, Pasadena, CA 91125, USA; Department of Chemistry, College of Chemistry, University of California, Berkeley, Berkeley, CA 94720, USA; Department of Electrical Engineering and Computer Sciences, University of California, Berkeley, Berkeley, CA 94720, USA; Division of Molecular Biophysics and Integrated Bioimaging, Lawrence Berkeley National Laboratory, Berkeley, CA 94720, USA; California Institute for Quantitative Biosciences (QB3), University of California, Berkeley, Berkeley, CA 94720, USA

**Author notes:** These authors contributed equally (Yikang Guan, Hengyu Li).

## Abstract

Cells maintain asymmetric electrochemical environments across selectively permeable plasma membranes. Ion channels on plasma membranes redistribute small ions, whereas large openings such as gasdermin pores allow both ions and proteins to cross the membrane, generating a multispecies transport regime. The transport of charged ions and proteins shapes electrochemical environments, leading to osmolarity changes, which in turn promotes water influx and cell swelling. Although biologists have observed the phenotypic progression from pore formation to cell swelling for decades, the molecular transport events underlying this progression remain elusive. To elucidate how pore formation drives downstream cellular effects, we establish quantitative multispecies transport frameworks to investigate gasdermin-perforated cells. We start by developing a reductionist model and benchmarking it against analytical Donnan predictions. We then explore three-dimensional geometries containing individual pores or pore clusters that form either simultaneously or sequentially. We reveal that ion-protein timescale separation generates non-canonical Donnan-like states, pore and cargo properties tune local transport, pore clustering modulates osmotic gradients, and sequential pore formation produces feedback-dependent dynamics. This work deepens our mechanistic understanding of cellular responses to membrane damage and informs the therapeutic design of pore-forming proteins with tunable cellular effects.

**Significance:** Large membrane pores mediate processes ranging from immune activation to inflammatory cell death, yet how pore opening produces cell swelling and rupture remains poorly understood. We develop a quantitative framework that follows ions and proteins as they move through gasdermin pores and reshape electrochemical conditions. Their unequal transport speeds create dynamic states not captured by classical equilibrium theory, while pore size, cargo size, pore clustering, and feedback-driven pore formation determine electrical and osmotic changes. These findings provide a physical explanation for how membrane damage drives water influx and swelling. More broadly, the framework establishes design principles for engineering pore-forming proteins whose downstream effects, including cell death, inflammation, and molecular delivery, can be tuned.

## Introduction

The plasma membrane separates the cytosol from the extracellular environment. Ion channels support ionic movement while retaining abundant charged macromolecules such as proteins within the cell.^1–3^ This selectivity changes when large pores disrupt the membrane. Such pores arise in diverse fundamental biological processes including immune cell hyperactivation and pyroptosis, a lytic form of cell death mediated by gasdermins. Following gasdermin pore formation, cells swell and may eventually undergo ninjurin-1 (NINJ1)-mediated plasma membrane rupture.^4–8^ While osmotic water influx has long served as the qualitative explanation for pore-induced cell lysis, the molecular transport events that alter the osmotic pressure remain challenging to describe quantitatively.

The Gibbs-Donnan effect offers a foundation for understanding how membrane pore formation drives osmotic water influx.^9–11^ In the classical Donnan scenario, immobile charged proteins in the cytosol lead to redistribution of permeant ions across the membrane, producing concentration differences, an electric potential, and an osmotic imbalance. By contrast, a gasdermin-perforated membrane presents a more dynamic situation. Gasdermin pores rapidly conduct small ions while also allowing the slower passage of proteins.^12–15^ Therefore, proteins act neither as a permanently fixed background, nor as a rapidly equilibrating solute. Instead, the system evolves across separated transport timescales, with ions rapidly responding to electrochemical gradients and proteins gradually altering the charge asymmetry. The Donnan effect therefore provides only a limiting reference for, or an early-time approximation, of multispecies transport in perforated cells.

While classical Donnan theory conveniently describes equilibrium states, it cannot resolve the evolving multispecies transport regime, in which spatial patterns of concentration, electric potential, and osmolarity emerge dynamically and vary with distance from pores. Understanding the evolution of electrochemical environments following pore formation necessitates a spatiotemporally resolved, quantitative description of multispecies transport.

The Poisson-Nernst-Planck equations, with the continuity, Nernst-Planck, and Poisson equations as the three coupled components, provide a rigorous framework for investigating ion transport. In this study, drawing inspiration from the application of these equations in ion transport,^16–21^ we explore the quantitative aspects of how gasdermin-perforated cells transport both ions and proteins. To do so, we integrate the distinct mobilities, sizes, and charges of ions and proteins to resolve their coupled transmembrane transport, with the goal of determining how electrochemical environments evolve and produce the osmotic driving force responsible for water influx following cell permeabilization by large membrane pores.

## Methods

### Multispecies transport framework

We developed one-dimensional and three-dimensional models to investigate how ion and protein transport reshape electrochemical environments following gasdermin-mediated membrane permeabilization (Figure 1A). Both models couple mass conservation, diffusion, electromigration, and electrostatics in cytosolic and extracellular aqueous compartments separated by a 4 nm membrane. The transported species comprise Na^+^, K^+^, Ca^2+^, Cl^−^, a negatively charged protein, a charged auxiliary species X, and a neutral osmolyte Y. The auxiliary species establish initial charge and osmotic balance. We calculated solute redistribution and the resulting electrical and osmotic responses in fixed computational domains, while allowing pore accessibility to change during sequential pore formation.

**Figure 1.**
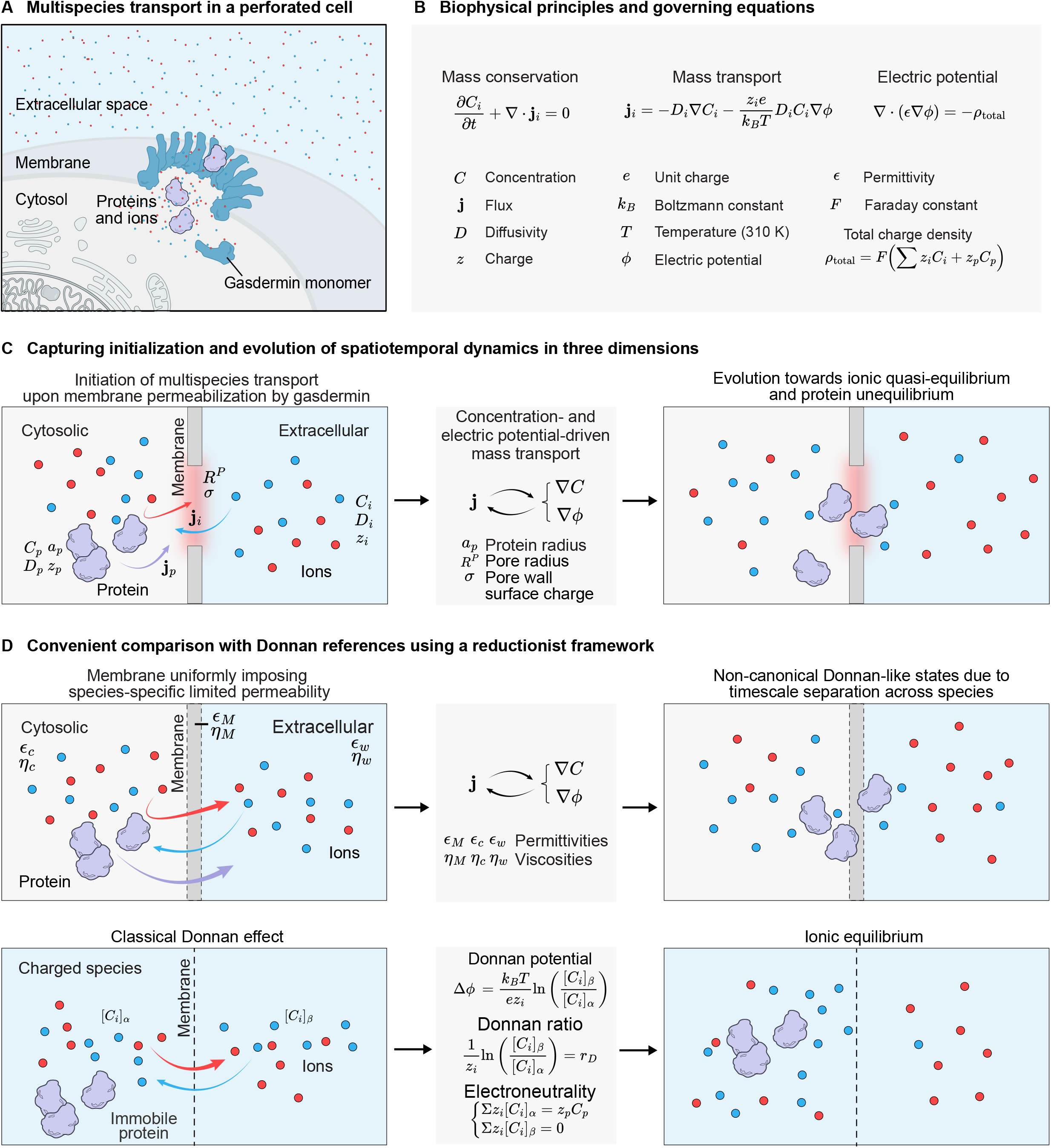
Quantitative frameworks capture multispecies transport in perforated cells. (A) Gasdermin monomers oligomerize into membrane pores that conduct both small ions and proteins between the cytosol and extracellular space. (B) The governing equations cover mass conservation, ion and protein flux by diffusion and electromigration, and evolution of the electric potential. (C) As ions and proteins cross the membrane through discrete gasdermin pores, the system relaxes toward a quasi-equilibrium of ions and protein redistribution. (D) Replacing discretely perforated membrane with a uniformly permeable membrane provides a far-field representation for convenient comparison with Donnan equilibrium.

For a species *i* with molar concentration *C*_*i*_ and flux *J*_*i*_, mass conservation requires

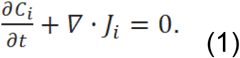

The conventional Nernst–Planck equation describes ionic transport through diffusion and electromigration:

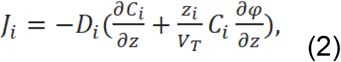

where *D*_*i*_ denotes diffusivity, *z*_*i*_ ionic valence, *e* elementary charge, *k*_*B*_ Boltzmann’s constant, and *T* = 310 *K* is temperature. The concentration gradient ∇*C*_*i*_ drives diffusion, whereas the electric potential gradient ∇ϕ drives electromigration. We determine the electric potential ϕ self-consistently from Poisson’s equation:

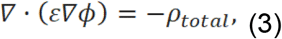

where *ε* is local permittivity. Both mobile-species and charged auxiliary species X contribute to the total charge, while neutral osmolyte Y contributes no charge. Prescribed fixed and pore-wall charges are added to total charge density where implemented. The total particle concentration determines osmotic pressure through van’t Hoff relation Π = *RTΣ*_*i*_C_*i*_, including proteins and auxiliary solutes.

### Reductionist model and Donnan reference

The reductionist one-dimensional model represents a perforated membrane as a uniformly permeable planar layer normal to the transport coordinate (Figure 1D). Regional permittivities *ε*_*c*_, *ε*_*M*_, and *ε*_*w*_ determine the electrostatic response, while viscosities *η*_*c*_, *η*_*M*_, and *η*_*w*_ determine region-dependent diffusivities, with subscripts *c*, *M*, and *w* denoting cytosol, membrane, and extracellular water, respectively. We assigned physiological ion concentrations and species-specific diffusivities, and calculated protein diffusivity using the Stokes–Einstein relation with hydrodynamic radius *R*_*H*_. The membrane permeability parameter *α*_*block*_ controls protein passage through the homogenized membrane layer.

The original 1D protein flux combines diffusion with separately specified electrophoretic mobility:

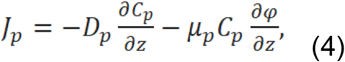

where *D*_*p*_ denotes protein diffusivity and *μ*_*p*_ denotes signed protein electrophoretic mobility. We estimated protein electrophoretic mobility using the Hückel expression with a Henry correction *f*_*H*_(*K a*_*p*_), where *a*_*p*_ is protein radius and *K* is inverse Debye length. The zeta potential ζ_*p*_ is estimated from protein charge and hydrodynamic radius. Regional permittivity and viscosity additionally enter the original mobility calculation. Consequently, the 1D protein mobility is a phenomenological electrokinetic approximation that does not strictly satisfy the conventional Nernst–Einstein relation. Its implementation is detailed in the Supporting Information.

We initialized the cytosolic and extracellular compartments at electroneutrality and equal osmolarity, then redistributed ions near the membrane to establish a resting potential of approximately −75 *mV*. We compared membrane-permeant proteins with effectively blocked proteins to distinguish the consequences of protein leakage.

Classical Donnan equilibrium provides a reference for the limiting case in which charged proteins remain confined to the cytosol while small solutes can redistribute (Figure 1D). The subscripts *α* and *β* denote the cytosolic and extracellular compartments, respectively. Equality of the electrochemical potentials of a permeant ion gives the Donnan potential:

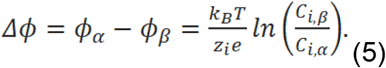

The Donnan partition ratio *r*_*D*_ is defined by

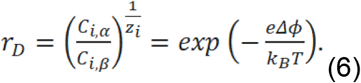

The retained protein charge *Z*_*p*_ *C*_*p*_ enters the electroneutrality condition, while permeant ions partition according to Eq. (6). This Donnan reference assesses the consequences of protein mobility and ion–protein timescale separation.

### Pore geometry and protein transport in three dimensions

The three-dimensional model explicitly resolves individual cylindrical gasdermin pores through an otherwise impermeable lipid membrane (Figure 1C). The default computational domain spans 1 *μ*m in each lateral direction and extends 1 *μ*m into each aqueous compartment. A nonuniform Cartesian mesh provides finer resolution near the membrane and pore centers. The symbols *R*^*P*^, *a*_*p*_, and *σ* denote pore radius, protein radius, and pore-wall surface charge density, respectively. The ion parameters *C*_*i*_, *D*_*i*_, and *Z*_*i*_, and protein parameters *C*_*p*_, *D*_*p*_, and *Z*_*p*_, describe their respective concentrations, diffusivities, and valences.

We excluded protein centers from the lipid membrane and from regions that cannot accommodate their finite radius. Within a cylindrical pore, the accessible lateral radius is *max*(*R*^*P*^, *a*_*p*_,0). We varied pore geometry, number, and spacing to examine individual pores and pore clusters. Surface charge is represented as a configurable parameter, with its numerical implementation described in the Supporting Information.

The default protein species has *C*_*p*_ = 1*mM*, *Z*_*p*_= −10, and *a*_*p*_ = 5*nm*. Protein-size dependence enters through Stokes–Einstein-based diffusivity and geometric accessibility.

We calculated the local occupied protein volume fraction *φ* by spatially averaging protein number density over spherical neighborhoods. A concentration-dependent crowding factor reduces both ion and protein transport:

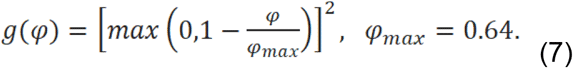

With thermal voltage *V*_*T*_ = *k*_*B*_ *T* / *e*, protein transport in the three-dimensional model follows

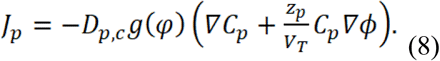

The same diffusivity and crowding factor enter the diffusion and electromigration terms, preserving the conventional Nernst–Einstein relation. Species-specific accessibility masks further restrict transport to permitted aqueous regions. Unlike the 1D formulation, the 3D protein flux does not include a Henry-function correction or an additional permittivity multiplier on electrophoretic mobility.

### Sequential pore formation

We compared simultaneous pore opening with sequential opening at prescribed times or in response to cumulative outward K^+^ transport. In the feedback condition, we integrated positive outward K^+^ molar flow across the membrane plane. Reaching successive threshold increments triggers additional pores, up to the configured maximum. This phenomenological rule connects potassium efflux to pore formation without explicitly modeling upstream receptor activation, inflammasome assembly, proteolysis, or gasdermin oligomerization.

### Numerical solution and analysis

The one-dimensional model evaluates ionic fluxes using Scharfetter–Gummel discretization and protein fluxes using discrete diffusion and centered electrophoretic drift. A tridiagonal Poisson solver updates the electric potential as concentrations evolve, while adaptive Runge–Kutta integration in SciPy advances the transport equations. We quantified fractional relaxation by normalizing each concentration trace to its net change over the observation window. The first crossings of 10%, 50%, and 90% define the corresponding relaxation times. The integral T_int_ of the remaining fractional change summarizes the cumulative relaxation lag.

The three-dimensional model uses a finite-volume discretization on a nonuniform mesh and advances transport with explicit Euler steps and Scharfetter–Gummel fluxes. A preconditioned conjugate-gradient solver updates the electric potential, and CuPy implements the arrays and transport kernels on an NVIDIA GPU.

Across the simulations, we tracked concentration profiles, membrane potential, osmolarity differences, protein leakage, and membrane current. The relaxation measures characterize dynamics within the observation window and do not establish convergence to equilibrium.

## Results

### Quantitative frameworks capture multispecies transport in perforated cells

Large membrane openings generate a multispecies transport regime in which ions and proteins cross the same membrane simultaneously. Gasdermin represents a biologically significant system to explore multispecies transport (Figure 1A). Upon activation by pathogenic signals or sterile dangers, gasdermin monomers assemble into pores with an inner diameter of around 215 Å, allowing the transport of both ions (e.g. Na^+^, K^+^, Ca^2+^, Cl^−^, etc.) and proteins (e.g. IL-1*β*, IL-18, HMGB1, cytochrome c, etc.).^22–24^ These species differ substantially in mobility, with ions rapidly redistributing whereas proteins moving more slowly.

To quantitatively depict multispecies transport, we integrated mass conservation, transport by diffusion and electromigration, and electric potential evolution into a unified biophysical framework (Figure 1B). Regarding mass conservation, the equation of continuity relates the change in concentration of each charged species to its mass flux.^25–27^ To describe the mass flux of ions, we used the Nernst-Planck equation, which accounts for both diffusion along concentration gradients and electrostatic drift along electric potential gradients.^28–30^ For proteins, we modified the Nernst-Planck equation to incorporate dependence on protein properties including size and charge. Because the redistribution of both ions and proteins alters the local electric field that in turn modulates their transport, we couple the continuity and Nernst-Plank equations to the Poisson equation, which relates the electric potential to charge density.^31–33^ These equations together provide a coherent quantitative description of multispecies transport spanning the cytosolic, membraneous, and extracellular regions with distinct permittivities and viscosities (Figure 1C). Within these regions, we initialize each species at defined intracellular and extracellular concentrations and resolve its redistribution over time following membrane permeabilization by gadsermin.

Before fully executing the spatiotemporally resolved framework, we simplified the geometry by replacing membrane pores with a membrane that uniformly imposes limited, species-specific permeability, representing the far-field view of a membrane containing many pores (Figure 1D). This geometry reduction offers three advantages. First, it substantially reduces the computational cost of the full three-dimensional simulations that explicitly resolve pore-containing and pore-free membrane regions. Second, it enables convenient validation of the continuity and Poisson-Nernst-Planck framework by testing whether it qualitatively reproduces the Donnan equilibrium that occurs in the context of protein immobility. Third, it allows us to quantify how multispecies transport deviates from Donnan equilibrium references when intracellular proteins can move across the membrane (Figure S1). This reductionist framework establishes a rigorous foundation for characterizing the emergence of evolving electrochemical environments in perforated cells.

### Non-canonical Donnan-like states arise from timescale separation

Having established the governing equations and simplified geometry, we studied the dynamic transport of ionic and protein cargoes following membrane permeabilization and compared the temporal profiles with analytical Donnan predictions (Figure 2A). To reflect biological scenarios, we used physiological intracellular and extracellular ionic concentrations while assigning each ion its aqueous diffusivity to preserve mobility differences. For simplicity, we represented proteins as a single mobile species carrying a charge of −10 e and a radius of 25 Å, consistent with the properties of a typical proteome.^34–36^ To enforce initial electroneutrality, we incorporated an additional unnamed mobile species that effectively accounts for the net contribution of all other ions and small solutes, including nucleotides and saccharides.

**Figure 2.**
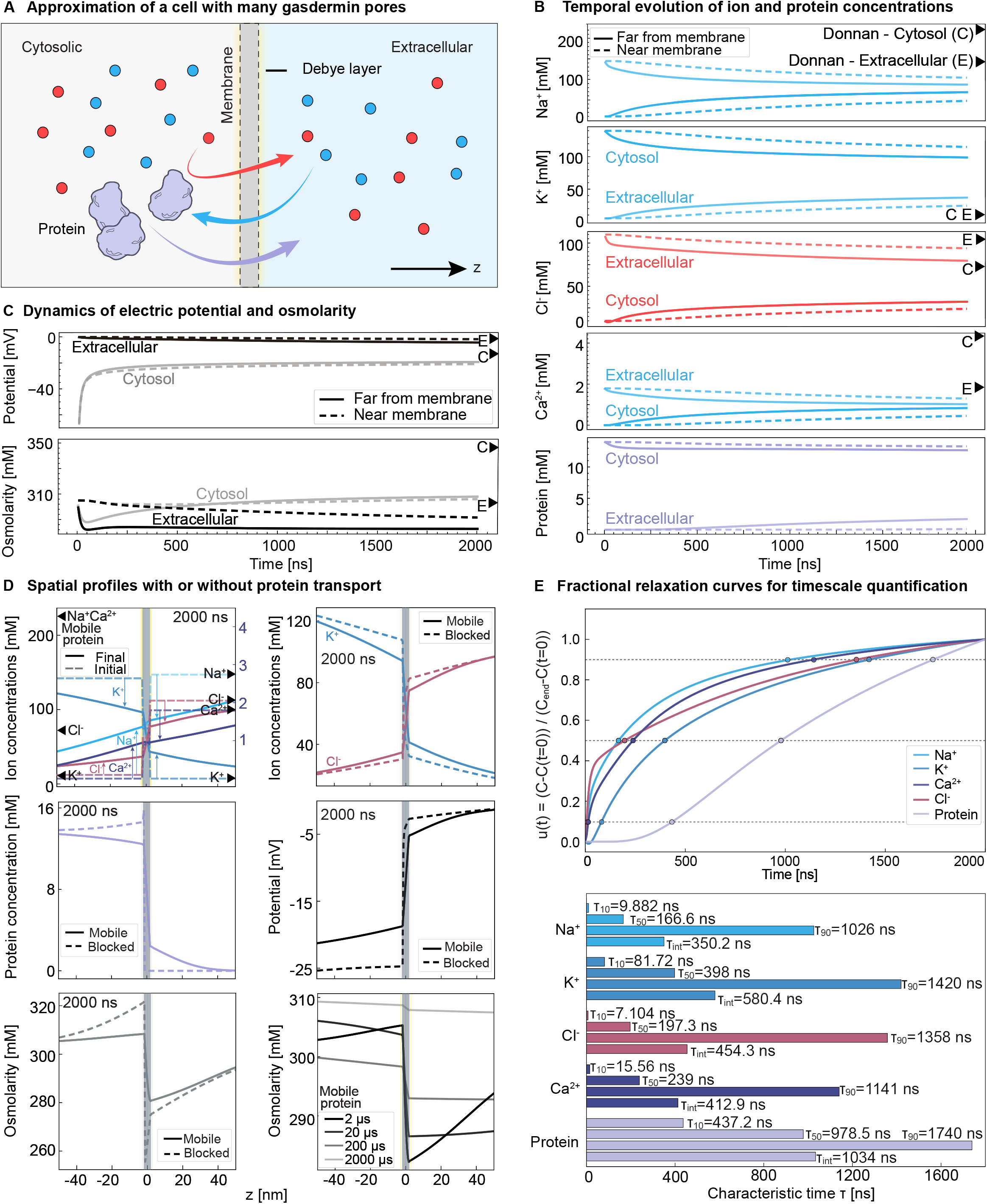
Non-canonical Donnan-like states arise from timescale separation. (A) A membrane separates cytosolic and extracellular compartments that contain ion and protein species with physiological diffusivities, sizes, charges, and concentrations. (B) Ion concentrations near and far from the membrane rapidly approach, but do not converge with, Donnan references, while protein concentrations change relatively modestly. (C) Cytosolic electrical potential initially exhibits a rapid increase and then gradually becomes less negative, while cytosolic osmolarity rises after a transient decrease. (D) Mobile proteins attenuate ion redistribution, narrow the transmembrane potential difference, and reduce osmotic imbalance relative to immobile proteins. (E) Normalized fractional-relaxation curves reveal a separation of transport timescales, with ions approaching unity more rapidly than proteins.

Tracking concentrations at representative positions located near (around 10 nm) and far (40 nm) from the membrane on both the cytosolic and extracellular sides, we revealed distinct temporal evolution of ion and protein transport (Figure 2B). Ions rapidly narrowed their transmembrane gradients within 2000 ns, whereas protein concentrations changed modestly. Near-membrane positions responded earlier and more strongly than membrane-distant regions, reflecting the local origin of transport. Of note, ion concentrations approached, but did not converge to, the Donnan reference values because proteins continued to leak through the pore. This ongoing protein transport maintained a non-canonical Donnan-like regime in which the ionic distributions remained dependent on boundary conditions and system parameters. Blocking protein transport restored the Donnan equilibrium, whereas protein mobility produced a balanced distribution of all species over extended time (Figure S2A and S2B).

The concentration changes of ions and proteins drove the coupled evolution of electric potential and osmotic pressure. The cytosolic potential became progressively less negative as cations influx dominated the total ion flux and negatively charged proteins exited the cell, while cytosolic osmolarity changed non-monotonically (Figure 2C). During the first 50-100 ns, electrostatic forces depleted co-ions near the membrane faster than diffusion replenished counter-ions from the bulk, transiently lowering osmolarity. Diffusion then dominated over the following hundreds to thousands of nanoseconds, increasing the transmembrane osmolarity difference. Like their temporal patterns, the spatial profiles of concentration, electric potential, and osmolarity depend on protein mobility (Figure 2D). Across the membrane, ion concentrations transition from their initial values toward, but not fully to, the classical Donnan predictions. Compared with impermeant proteins, mobile proteins attenuate ion redistribution, narrow the transmembrane potential difference, and reduce the osmotic imbalance across the membrane.

To further rationalize the spatiotemporal observations, we quantified ion and protein relaxation using a model-free fractional-relaxation metric, u(t), tracking the time each species required to reach 10%, 50%, and 90% of its total concentration change (Figure 2E). Ions consistently reached half-relaxation within a few hundred nanoseconds, whereas the protein required nearly 1000 ns to reach the same fractional threshold, confirming a clear separation in relaxation speed without relying on parametric fit. The persistent lag drives the system into a non-canonical Donnan-like state, and because proteins continue to leak while ions rapidly re-equilibrate, ions relax toward a moving target rather than a fixed equilibrium. Therefore, timescale separation underlies the evolving electrochemical environments of a perforated cell.

### Pore and cargo properties tune spatiotemporal transport patterns

Moving beyond the simplified geometry, we next resolved multispecies transport around a single pore using a full three-dimensional framework (Figure 3A). To reduce computational cost, we analyzed transport over 1000 ns, which was sufficient to reveal the spatiotemporal evolution of ion concentrations and electric potential (Figure 3B and S3). Near the pore, cytosolic osmolarity increased while extracellular osmolarity decreased due to Donnan-like redistribution of ions and proteins. Immediately adjacent to the pore, however, the osmolarity pattern reversed because the resting membrane potential and intact Debye layer remained largely preserved.

**Figure 3.**
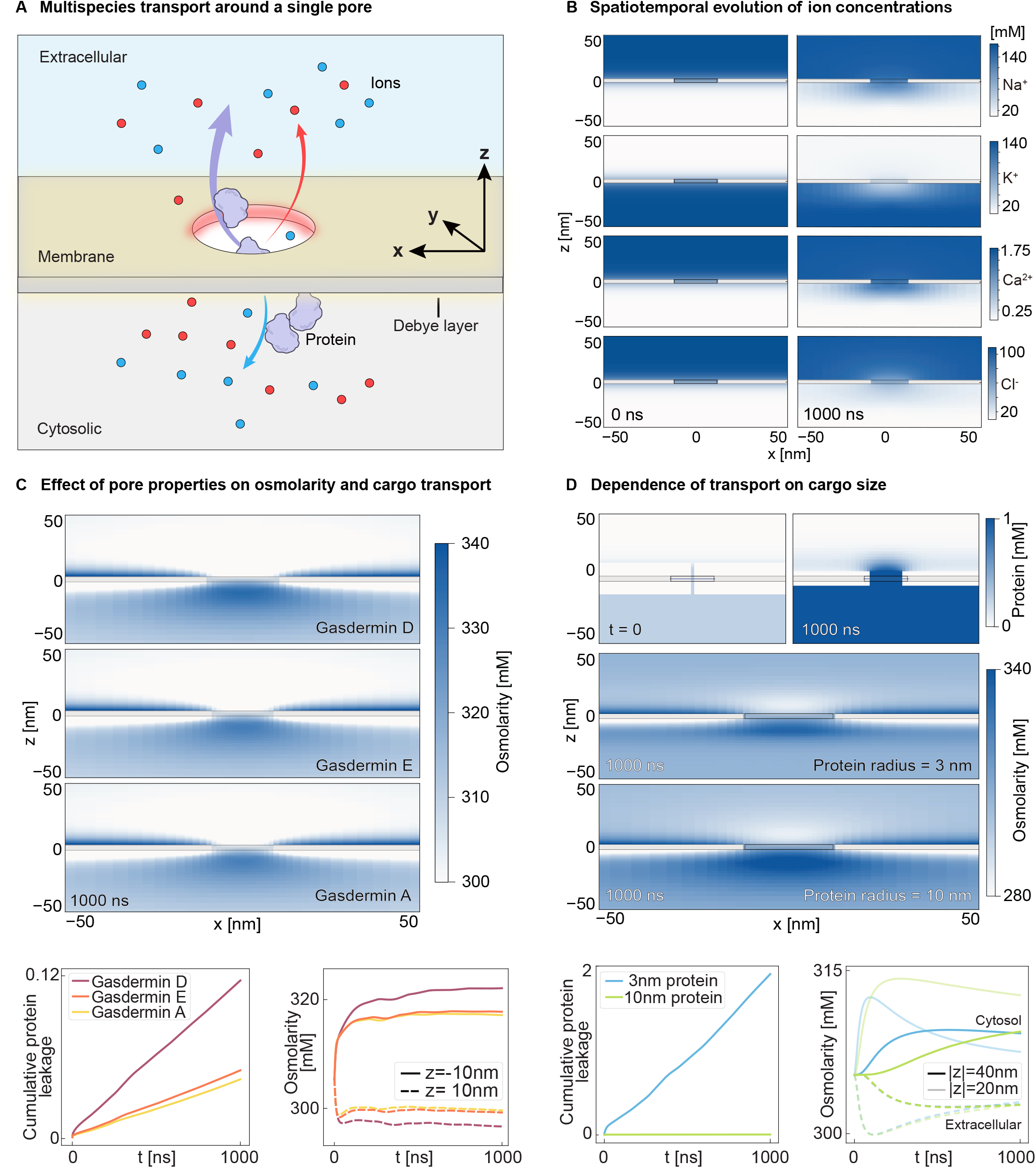
Pore and cargo properties tune spatiotemporal transport patterns. (A) A full three-dimensional framework resolves multispecies transport around a single gasdermin pore embedded in a membrane with Debye layers. (B) Ion flux breaks Debye layers around the pore and rapidly begins to decrease ion concentration gaps across the membrane. (C) Three representative gasdermin pores demonstrate that pore size and charge density influence surrounding electrochemical environments during multispecies transport. (D) Larger proteins undergo more entrance accumulation, thereby reducing permeation, increasing the cytosolic osmolarity near the membrane and decreasing it farther away.

The pore-scale transport patterns above suggested that properties of the pore itself may regulate multispecies transport. We therefore evaluated how pore size and charge density influence surrounding electrochemical environments using representative gasdermin pores (Figure 3C). Quantification of visually modest differences in the spatial profiles revealed progressively greater protein leakage and intracellular osmolarity with increasing pore size and charge density. These observations are consistent with the underlying transport mechanisms, where larger pores reduce steric hindrance for protein transport, whereas more negatively charged pores impede the transport of negatively charged proteins.^37–42^ Across all pore sizes and charges, intracellular osmolarity is consistently higher than extracellular osmolarity, with the largest transmembrane osmotic difference occurring near the pore and decreasing with distance from the membrane, suggesting that water influx may preferentially occur around the pore.

After examining pore properties, we wondered whether the size of protein cargoes influences electrochemical environments during multispecies transport (Figure 3D). We represented protein-size dependence through the inverse-radius dependence of the effective Stokes–Einstein diffusivity and geometric constraints on protein accessibility. Protein centers must remain at least one protein radius from the pore wall, progressively narrowing the accessible opening as protein size increases. While proteins initially remained nearly homogeneous throughout the cytosol, they accumulated within 1000 ns near the cytosolic pore entrance. This entrance accumulation effect arises because concentration gradients drive protein traffic toward the pore, while steric hindrance restricts passage through the pore. Quantification revealed that increasing the protein radius from 3 to 10 nm nearly abolished protein leakage, even though a 10 nm protein remains smaller than the approximately 215 Å pore diameter, consistent with prior experimental observations.^43–45^ The retention of intracellular proteins increased cytosolic osmolarity and enlarged the osmotic difference near the membrane, while reducing cytosolic osmolarity farther from the pore because the proteins accumulated locally rather than redistributing throughout the cytosol.

Together, these results extend the non-canonical Donnan-like transport framework from the reductionist model to pore-scale phenomena in three-dimensional geometry, demonstrating that both pore architecture and cargo properties leave distinct signatures on surrounding electrochemical environments.

### Pore density and cluster area modulate electrochemical outputs

Single-pore transport in three-dimensional geometry isolates the transport behavior of an individual pore but cannot capture interactions among pores. Confocal microscopy revealed that gasdermin pores assemble into puncta large enough to encompass clusters of pores,^46–48^ suggesting that the spatial organization of pores may reshape surrounding electrochemical environments. To investigate this possibility, we examined how pore density (number of pores per unit cluster area) and cluster area regulate multispecies transport. More specifically, we compared a single pore, two 16-pore (4×4) clusters with different pore densities, and a 64-pore (8×8) cluster with a larger cluster area (Figure 4A).

**Figure 4.**
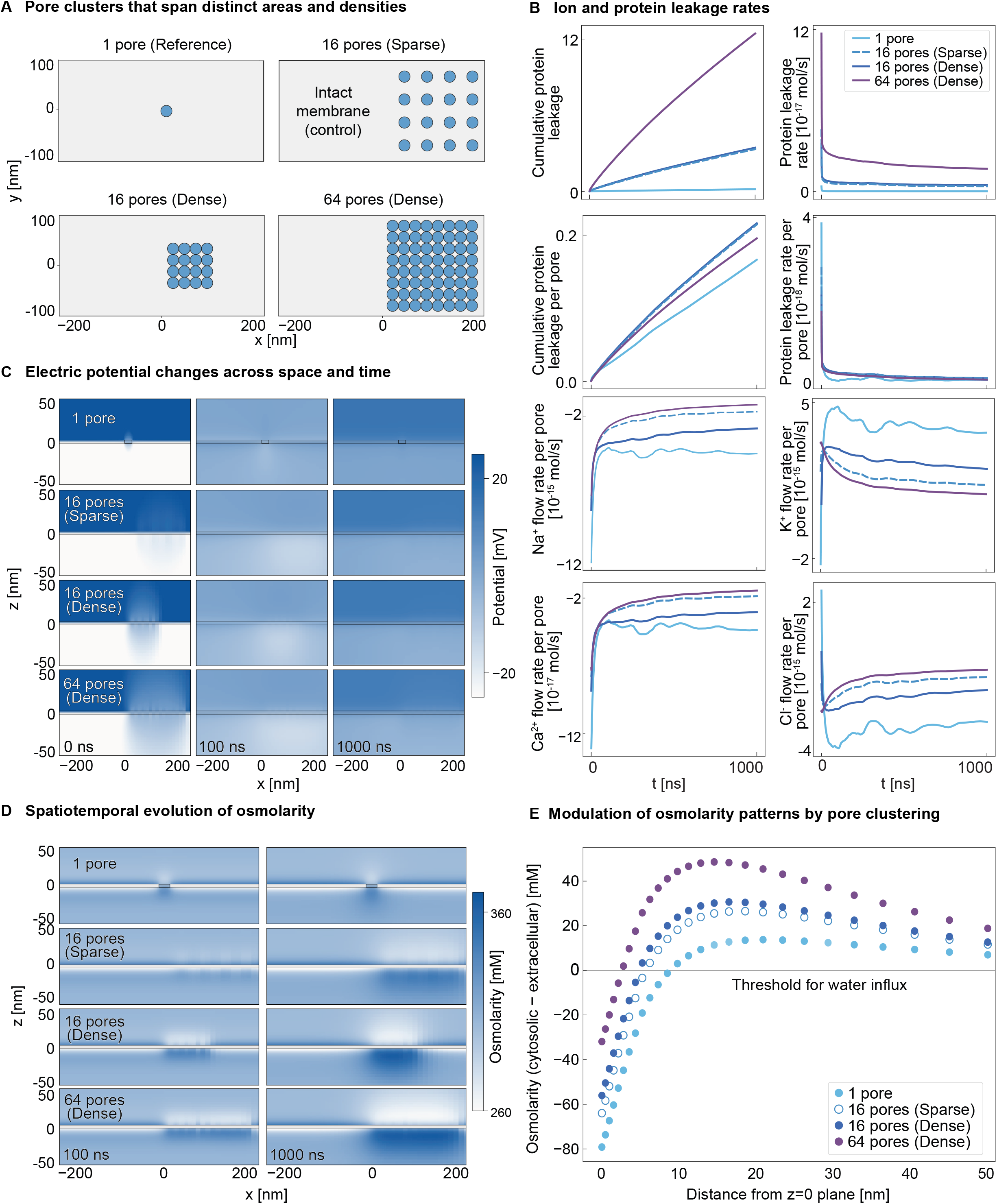
Pore density and cluster area modulate electrochemical outputs. (A) Four spatial configurations, a single pore, sparse and dense 16-pore (4×4) clusters, and a dense 64-pore (8×8) cluster illustrate the effects of pore density and cluster area. (B) Cumulative protein leakage scales with cluster area, per-pore protein efflux converges across clusters, whereas per-pore ion flux decreases with pore density and cluster area. (C) Larger clusters and higher densities initially generate stronger potentials, but lateral ion movement quickly homogenizes them across all configurations. (D) Osmolarity maps show that higher pore density and larger cluster area progressively enhance and spatially expand elevated cytosolic osmolarity over time. (E) The cytosolic-extracellular osmolarity difference approaches the threshold for osmotic water influx at a shorter distance from the membrane as pore density or cluster area increases.

The results reveal that cluster area and pore density differentially shape protein and ion transport. Cumulative protein leakage increased with cluster area, which reflects the greater total number of pores, and remained agnostic of pore density, whereas per-pore protein efflux converged across all spatial configurations, indicating that larger clusters enhance total protein release primarily by increasing pore number rather than the transport efficiency of individual pores (Figure 4B). This behavior suggests that the relatively low abundance of intracellular proteins allows individual pores to sample the protein pool largely independently. In contrast, cumulative ion flux increased with either higher pore density or larger cluster area, while per-pore ion flux decreased under both conditions, likely due to competition among neighboring pores for the same local ionic reservoir, indicating that pore clustering amplifies ionic transport beyond simply scaling the single-pore response (Figure 4C).

To better elucidate how pore clustering shapes membrane-scale electrochemical environments, we next examined its effects on electric potential and osmolarity. While larger cluster areas and higher pore densities initially generated stronger and more spatially extensive electric potentials, lateral ion movement progressively homogenized the potentials, with all spatial configurations becoming nearly identical by 1000 ns (Figure 4D). In contrast, osmolarity exhibited a distinct response. After 100 ns, the low-density 16-pore cluster modestly, whereas the high-density 16-pore cluster dramatically, enhanced cytosolic osmolarity in comparison with single-pore reference. This enhancement became more pronounced over time, indicating that spatial proximity of pores amplifies osmolarity (Figure 4E). Comparing the 16- and 64-pore clusters with the same pore density further showed that increasing cluster area expanded the membrane region exhibiting elevated cytosolic osmolarity. Thus, while pore density and cluster area primarily modulate the transient kinetics of electric potential, they strongly shape the magnitude and spatial extent of osmolarity over time.

The prominent effect of pore clustering on osmolarity raised the question of why cells assemble pores into clusters. We hypothesized that pore clustering generates cytosolic-high, extracellular-low osmolarity patterns that drive water influx. To test this hypothesis, we quantified the cytosolic-to-extracellular osmotic pressure difference as a function of distance from the membrane across different pore configurations (Figure 4E and S4). Increasing either pore density or cluster area brought a given osmotic pressure difference closer to the membrane. Consequently, if water influx necessitates a threshold osmotic pressure difference, pore clustering would effectively lower the threshold for localized water influx. These findings reveal that the spatial organization of pores, beyond the properties of an individual pore, functions as a critical determinant of electrochemical environments in perforated cells.

### Sequential pore formation generates feedback-dependent dynamics

The preceding findings provided insights into multispecies transport in both simplified and full three-dimensional geometries, considering individual pores as well as pore clusters that form simultaneously. However, membrane perforation is fundamentally a dynamic and sequential process. For instance, during pyroptosis, K^+^ efflux through existing gasdermin pores activates inflammasomes, which in turn activate caspases that proteolytically activate additional gasdermins, forming a positive feedback loop that promotes further pore formation.^49–51^ Unlike the static pore clusters, this signaling cascade involves multiple diffusible intermediates, allowing new pores to form throughout the plasma membrane rather than immediately adjacent to existing pores. To complete this study, we investigated how feedback-driven sequential pore formation shapes electrochemical environments (Figure 5A).

**Figure 5.**
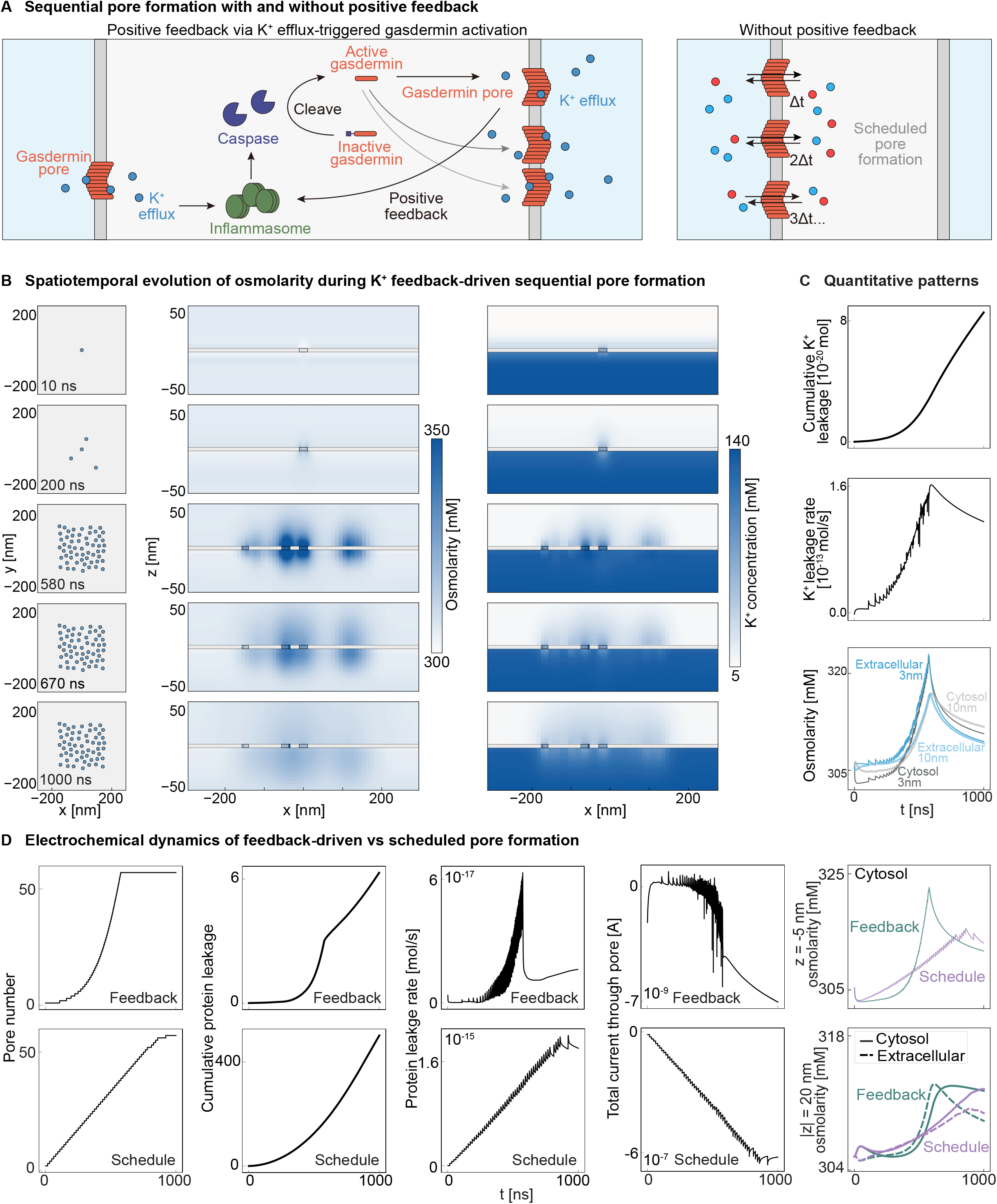
Sequential pore formation generates feedback-dependent dynamics. (A) K^+^ efflux activates gasdermin through multiple signaling molecules, unlike a reference scenario with sequential signaling-independent pore insertion. (B) As pores accumulate sequentially, extracellular osmolarity near the membrane transiently exceeds cytosolic osmolarity due to rapid K^+^ spikes. (C) During the early phase, K^+^ efflux and osmolarity increase exponentially, consistent with characteristic dynamics driven by positive feedback. (D) Positive feedback lets pore formation outpace electrochemical relaxation, causing non-monotonic excursions and delayed convergence to Donnan-like quasi-steady states.

As pores accumulated sequentially under K^+^ efflux-mediated positive feedback, each newly inserted pore locally generated the characteristic cytosolic-high, extracellular-low osmolarity pattern (Figure 5B and S5). As pore number increased, however, extracellular osmolarity near the membrane transiently exceeded cytosolic osmolarity despite membrane-distant regions remaining cytosolic-high. This transient reversal arose because K^+^ dominated the initial transport response. Its high intracellular abundance relative to Ca^2+^ and rapid diffusivity relative to Na^+^ produced an immediate extracellular K^+^ spike following each pore insertion, whereas redistribution of the remaining ionic species occurred more gradually, an electrochemical signature consistent with the K^+^-mediated feedback mechanism itself. After the pore insertion cascade concluded, the electrochemical environment progressively relaxed toward the non-canonical Donnan-like quasi-steady state as the remaining ions redistributed and proteins accumulated near pore entrances. Thus, although sequential pore formation ultimately converged to the same cytosolic-high, extracellular-low osmolarity state observed for single pores and static pore clusters, it traversed multiple transient, non-monotonic intermediate states. The transient reversals in the osmolarity gradient suggest that water influx may occur intermittently rather than continuously during membrane perforation, and if so, would explain experimentally observed progressive cell swelling preceding pyroptotic cell lysis.^52–56^

Further quantification of the spatiotemporal dynamics deepened our understanding of how positive feedback shapes pore formation and multispecies transport (Figure 5C). Pore formation and its downstream consequences, including protein leakage and the increase in near-membrane cytosolic osmolarity, initially followed exponential trajectories characteristic of positive feedback. Once pore insertion events concluded, the system gradually relaxed toward noncanonical Donnan-like quasi-steady states. Average osmolarity near the membrane remained higher outside the cell throughout the insertion loop and reversed only after pore formation ended. By contrast, regions farther from the membrane became cytosolic-high earlier during the loop because slower pore insertion allowed greater relaxation between successive insertion events. Lastly, to isolate the functional role of positive feedback, we finally investigated sequential pore insertion without it (Figure 5D). Unlike the exponential dynamics produced by feedback, which allowed new pore formation to outpace electrochemical relaxation and thereby delayed convergence toward the noncanonical Donnan-like quasi-steady state, constant-rate pore insertion generated approximately linear transport dynamics, allowing electrochemical relaxation to proceed concurrently with pore formation.

In summary, sequential pore formation with positive feedback enriches the dynamics of electrochemical environments by generating transient, non-monotonic behaviors and delaying relaxation, revealing a rich quantitative landscape of transport phenomena in perforated cells.

## Discussion

In this study, we established quantitative frameworks with complementary geometries to investigate multispecies transport in perforated cells, connecting cargo properties, pore properties, pore spatial organization, and the temporal sequence of pore formation with electrochemical dynamics across molecular, spatial, and temporal scales.

Our findings provide a quantitative foundation for understanding cellular responses to membrane damage, a process that underlies diverse physiological and pathological contexts. By resolving the spatiotemporal evolution of ions, proteins, electric potential, and osmolarity, we establish a physical mechanism linking membrane perforation to osmotically driven water influx and cell swelling, which may mechanically promote NINJ1-mediated plasma membrane rupture. This terminal rupture is distinct from loss of viability, which can occur earlier, including through gasdermin-mediated mitochondrial damage, and instead enables broad release of intracellular contents that increases exposure of the dying cell to the immune system.

Membrane perforation in cells encompasses greater mechanistic complexity than the scenarios we describe here. Membrane lipid composition, curvature, geometry, and subcellular localization all influence transport, motivating future integration of our frameworks with in-depth molecular dynamics simulations and experimental measurements. Likewise, small molecules and macromolecules exhibit diverse biochemical identities both intracellularly and extracellularly, including nucleic acids, saccharides, and metabolites that span broad ranges of concentration, size, charge, and diffusivity. Incorporating these additional species and their distinct features into multispecies transport will likely uncover more sophisticated spatiotemporal dynamics.

Finally, the ability to quantitatively describe electrochemical environments will guide the rational design of pore-forming proteins with customizable cellular effects. Current therapeutic strategies based on pore-forming proteins are efficient at triggering membrane perforation while limited in the ability to explicitly program downstream cellular responses.^57–60^ Our findings indicate that ionic redistribution, protein transport, membrane potential, and osmotic gradients are tunable during membrane perforation. Because membrane perforation broadly mediates cell death, inflammation, and intracellular cargo delivery,^61–64^ making its biochemical and cellular consequences as programmable as pore formation itself will substantially broaden the therapeutic utility of pore-forming proteins.

## Data and code availability

All code and data supporting the findings in the main and supplementary figures will be deposited in Github (https://github.com/c1462881/Multispecies-transport-model) before publication of this paper. Any additional information required to reanalyze the data reported here is available from the lead contact upon request.

## Acknowledgments

We thank Kranthi Mandadapu, Karthik Shekhar, Sirui Ning, Babak Vajdi Hokmabad, and all members of the Shiyu Xia lab for insightful discussions; Inna Marie Strazhnik for graphical design; and Lauren Hughes, Kris Stangl, Sigrid Allen, and Marlyn Grant for administrative support. This work was supported by the Burroughs Wellcome Fund Career Awards at the Scientific Interface, the Jane Coffin Childs Memorial Fund for Medical Research, and generous startup funding from the University of California, Berkeley.

## Author contributions

Y.G. performed computation, data analysis, and figure preparation. H.L. analyzed the data with Y.G. and identified directions for model refinement and further analysis. S.X. and Y.G. wrote the manuscript, with key comments and revisions from H.L., M.S., K.S., and Y.B.G. that improved the manuscript. S.X. conceived and supervised the project, with generous input from departmental colleagues.

## Declaration of generative AI and AI-assisted technologies in the manuscript preparation process

During the preparation of this work, the authors used Claude (Anthropic) and ChatGPT (OpenAI) to improve the readability of the manuscript. The authors reviewed and edited the output as needed and take full responsibility for the content of the published article.

