## Supplementary Information for "Multispecies transport shapes electrochemical environments in perforated cells"

### Supporting Information

### Methods

#### 1. General definitions and numerical conventions

The one-dimensional and three-dimensional implementations follow the conservation, Nernst–Planck, and Poisson framework introduced in Eqs. (1)–(3) of the main text. The coordinate normal to the membrane is $z$, with the cytosol at negative $z$ and the extracellular compartment at positive $z$. Positive flux denotes transport from cytosol to extracellular space.

The temperature is $T = 310$ K. We define the thermal voltage as $V_T = k_B T/e = RT/F$, where $R$ is the gas constant and $F = N_A e$ is Faraday's constant. Small-solute concentrations use units of mol m⁻³, which are numerically equal to mM. The 1D model expresses protein concentration as the molar concentration $C_p$, whereas the 3D model stores it as the number density $n_p$, with $C_p = n_p/N_A$. The calculations convert consistently between these representations when evaluating charge density and osmolarity.

The instantaneous charge distribution determines the electric potential. Permittivity varies spatially between aqueous and lipid regions. The sum of physiological ions, proteins, and auxiliary species X and Y determines the osmotic pressure. The computational geometries remain fixed, except for changes in aqueous accessibility when new pores are inserted in the 3D model. The model does not explicitly evolve solvent velocity, membrane deformation, or osmotic water flow.

#### 2. Reductionist one-dimensional model

##### 2.1 Geometry, species, and physical parameters

The reductionist model resolves transport along $z \in [-500{,}500]$ nm, using $N = 2000$ uniformly spaced grid points with spacing $\Delta z \approx 0.5003$ nm. The membrane occupies $|z| \leq 2$ nm and has a nominal thickness of 4 nm. The cytosol occupies $z < -2$ nm, and the extracellular solution occupies $z > 2$ nm. We analyze spatial profiles within $|z| \leq 50$ nm and monitor near- and far-membrane positions at approximately $z = \pm 10$ and $\pm 40$ nm.

The physiological species are $Na^{+}$, $K^{+}$, $Cl^{-}$, and $Ca^{2+}$. Initial cytosolic/extracellular concentrations are 10/145, 140/5, 10/110, and 0.0001/1.8 mM, respectively. The corresponding aqueous diffusion coefficients are 1.33, 1.96, 2.03, and $0.79 \times 10^{-9}$ m² s⁻¹.

The model protein has valence $z_p = -10$, physical radius $a_p = 2.5$ nm, and hydrodynamic radius $R_H = 1.3a_p = 3.25$ nm. Local electroneutrality determines its initial cytosolic concentration. Auxiliary species X is a charged, membrane-permeant balancing solute, while Y is a neutral, membrane-permeant osmolyte. Both have aqueous diffusivity $2.0 \times 10^{-9}$ m² s⁻¹.

The relative permittivities are 80 in aqueous regions and 2 in the membrane. The water, cytosolic, and membrane viscosities are $\eta_w = 0.691$, $\eta_c = 1.2$, and $\eta_M = 10$ mPa s, respectively.

##### 2.2 Regional diffusivities and protein mobility

The 1D model uses the conventional Nernst–Planck equation for physiological ions and the mobile auxiliary species. Ion diffusivities are reduced within the membrane according to the viscosity ratio:

$$D_{i,M} = D_{i,0} \frac{\eta_w}{\eta_M}. \quad \text{(S1)}$$

Outside the membrane, physiological ions retain their assigned aqueous diffusion coefficients.

Protein diffusivity in water follows the Stokes–Einstein relation:

$$D_{p,w} = \frac{k_B T}{6\pi\eta_w R_H}. \tag{S2}$$

The regional protein diffusivities are

$$D_{p,c} = \frac{\eta_w}{\eta_c} D_{p,w}, \qquad D_{p,M} = \alpha_{\text{block}} \frac{\eta_w}{\eta_M} D_{p,w}. \tag{S3}$$

Here, $\alpha_{\text{block}}$ controls the effective permeability of the homogenized membrane to proteins. The supplied default configuration uses $\alpha_{\text{block}} = 10^{-20}$, making protein transport through the membrane negligible. The permeant-protein condition uses $\alpha_{\text{block}} = 1$.

In the original 1D model, protein diffusion and electrophoretic drift are implemented separately, as expressed in Eq. (4). The Hückel expression, modified by Henry's function, gives the signed electrophoretic mobility:

$$\mu_p = \frac{2\varepsilon\zeta_p}{3\eta} f_H(\kappa a_p), \qquad \zeta_p = \frac{z_p e}{4\pi\varepsilon_w R_H}. \tag{S4}$$

The Henry correction is

$$f_H(\kappa a_p) = 1 + \frac{1}{2}\left[1 + \frac{2.5}{\kappa a_p(1+2e^{-\kappa a_p})}\right]^{-3}. \tag{S5}$$

Here, $\kappa$ is inverse Debye length, $\zeta_p$ is the estimated protein zeta potential, and $\varepsilon_w$ is aqueous permittivity. The mobility depends on the viscosity and permittivity of the corresponding region. Within the membrane, the calculation also includes $\alpha_{\text{block}}$. The supplied 1D implementation calculates the Henry factor from the initial extracellular Debye length and then holds it constant.

Combining Eqs. (S2)–(S5), the relationship between the original electrophoretic mobility and the local Stokes–Einstein diffusivity is

$$\frac{\mu_p}{z_p e D_p/(k_B T)} = \frac{\varepsilon}{\varepsilon_w} f_H(\kappa a_p). \tag{S6}$$

In aqueous regions, the ratio is $f_H(\kappa a_p)$; in the membrane, it is $(\varepsilon_M/\varepsilon_w) f_H(\kappa a_p)$. For the specified conditions, $\kappa a_p \approx 2.92$ and $f_H \approx 1.090$.

Thus, the original 1D protein electrophoretic mobility is a phenomenological electrokinetic coefficient that differs from the conventional Nernst–Einstein mobility based on $D_p$ and the nominal protein charge. The regional permittivity multiplier represents an additional modeling assumption independent of Henry's function. Both features are specific to the original reductionist formulation and are absent from the 3D protein flux.

##### 2.3 Initial electroneutrality and osmotic balance

The model initializes physiological ions at their prescribed cytosolic and extracellular concentrations and sets their membrane concentrations to zero. Protein initially occupies only the cytosolic region.

Local electroneutrality sets the cytosolic protein concentration:

$$C_{p,c}(0) = -\frac{\sum_i z_i C_{i,c}(0)}{z_p}. \tag{S7}$$

For the supplied ion concentrations, $C_{p,c}(0) = 14$ mM.

Initially, the charged auxiliary species X occupies only the extracellular compartment. Its charge sign and concentration neutralize the net charge of the physiological extracellular ions. Under the specified conditions, $z_X = -1$ and $C_{X,w}(0) = 43.6$ mM. The model adds neutral species Y to the compartment with the lower initial particle concentration, giving $C_{Y,c}(0) \approx 131.4$ mM and $C_{Y,w}(0) = 0$.

These conditions yield equal initial bulk osmolarities of approximately 305.2 mM, corresponding to an ideal osmotic pressure of approximately 787.1 kPa at 310 K. X and Y subsequently evolve as mobile species; they are not maintained as fixed background concentrations throughout the simulation.

Redistributing physiological ions within narrow slabs adjacent to the two membrane faces establishes the initial membrane potential. The extracellular Debye length is

$$\lambda_D = \left(\frac{\varepsilon_w k_B T}{e^2 N_A \sum_i z_i^2 C_{i,w}}\right)^{1/2}. \quad \text{(S8)}$$

For the specified conditions, $\lambda_D \approx 0.857$ nm. The slab thickness is set to $\max(\lambda_D, 1.5\Delta z)$. Weights scale with $z_i^2 \overline{C}_i$, where $\overline{C}_i$ denotes the mean initial concentration across the cytosolic and extracellular compartments. The shifts conserve each physiological species between the two slabs, while per-species clipping prevents negative donor concentrations.

Bracketing and bisection determine a scalar shift magnitude from the Poisson-resolved difference between the mean cytosolic and extracellular potentials. The target is $V_{\text{rest}} = -75$ mV, with an acceptance tolerance of 0.5 mV.

##### 2.4 Donnan equilibrium as a reference

Classical Donnan equilibrium provides a reference for the limiting case in which proteins remain intracellularly confined while smaller permeant species equilibrate electrochemically. The cytosolic and extracellular compartments correspond to $\alpha$ and $\beta$, respectively, as defined in the main text.

For an ideal permeant ion, equality of electrochemical potentials gives

$$RT\ln\left(\frac{C_{i,\alpha}}{C^\circ}\right) + z_i F \phi_\alpha = RT\ln\left(\frac{C_{i,\beta}}{C^\circ}\right) + z_i F \phi_\beta.$$

Consequently, the concentrations obey the partition relation $C_{i,\alpha} = C_{i,\beta} r_D^{z_i}$, with the Donnan potential and ratio defined in Eqs. (5) and (6).

The 1D implementation calculates a reservoir Donnan reference by holding the extracellular concentrations at their prescribed initial values and solving intracellular electroneutrality:

$$\textstyle\sum_{i \in \mathcal{M}} z_i\, C_{i,\beta} r_D^{z_i} + z_p C_{p,c} = 0, \quad \text{(S9)}$$

where $\mathcal{M}$ contains the physiological ions and charged auxiliary species X. The neutral osmolyte Y contributes no charge.

The resulting concentrations and potential difference are

$$C_{i,\alpha}^D = C_{i,\beta} r_D^{z_i}, \qquad \Delta\phi_D = -V_T \ln r_D. \quad \text{(S10)}$$

The Donnan osmotic pressure is calculated from the corresponding concentrations of all permeant species, retained protein, and neutral osmolyte. Because the extracellular concentrations are fixed in this calculation, the reference is a reservoir equilibrium and does not enforce the mass conservation of the finite simulation domain. It is therefore a thermodynamic comparison state, rather than a prediction of the finite-domain trajectory's terminal condition.

The familiar four-ion Donnan relations can also be recovered analytically when auxiliary ions are omitted. With $k = r_D^{-1}$, monovalent cations partition by $k$, monovalent anions by $1/k$, and divalent cations by $k^2$ when concentrations are expressed from the intracellular to extracellular side. This simplified construction illustrates the negative intracellular Donnan potential generated by trapped negative charge. For this study, Eq. (S9) provides the numerical reference and includes the charged auxiliary species.

##### 2.5 Spatial discretization and time integration

Harmonic averages of permittivity at grid faces discretize Poisson's equation. The scipy.linalg.solve_banded routine solves the resulting tridiagonal linear system. At the boundaries, the cytosolic end has zero potential gradient and the extracellular end has $\phi = 0$.

Physiological ions and auxiliary species are transported using the Scharfetter–Gummel face flux:

$$J_{i,f} = \frac{D_{i,f}}{\Delta z}\left[B(\psi_i)C_{i,L} - B(-\psi_i)C_{i,R}\right], \tag{S11}$$

where

$$\psi_i = \frac{z_i(\phi_R - \phi_L)}{V_T}, \qquad B(x) = \frac{x}{e^x - 1}. \tag{S12}$$

Here, $L$ and $R$ denote the adjacent grid nodes, and $D_{i,f}$ is the arithmetic mean of their diffusivities. A small-argument series evaluates the Bernoulli function for $|x| \leq 10^{-6}$. Elsewhere, the calculation clips its argument to $[-500,500]$.

In the original 1D protein implementation, the diffusion component uses Eq. (S11) with zero valence, followed by a centered electrophoretic drift term. Arithmetic averages of adjacent nodes define face mobility and protein concentration. A flux-scaling limiter and nonnegative concentration clipping suppress numerical undershoots.

The extracellular outer boundary holds physiological ions, X, and Y at their initial reservoir concentrations. The remaining transport boundaries use the implemented end-node derivative extension, in which the end-node time derivative is copied from the adjacent interior node. Protein does not have a fixed-concentration reservoir boundary.

The explicit RK45 method in scipy.integrate.solve_ivp integrates the semidiscrete equations with a relative tolerance of $10^{-4}$ and a componentwise absolute tolerance of $10^{-6}$ mM. The maximum integration step is

$$\Delta t_{\max} = \min\left(100\frac{\Delta z^2}{2D_{\max}}, \frac{T_{\text{sim}}}{50}\right), \tag{S13}$$

where $D_{\max}$ is the largest assigned physiological-ion aqueous diffusivity. The simulation duration $T_{\text{sim}}$ is specified in nanoseconds, and output times are uniformly sampled. The supplied default configuration uses $T_{\text{sim}} = 2{,}000{,}000$ ns and 300 output times; shorter analysis windows are also used for the early-time comparisons.

##### 2.6 Numerical diagnostics and relaxation analysis

At each saved time, the instantaneous charge density determines the electric potential, and the particle concentrations determine the osmotic pressure:

$$\Pi = RT\left(\sum_i C_i + C_p + C_X + C_Y\right), \tag{S14}$$

where the ionic summation excludes X and Y to avoid double counting. The model exports spatial concentration and potential profiles, local temporal measurements, and calculated osmotic pressures. We also check numerical mass conservation and minimum concentrations.

Both fitted and model-free measures characterize relaxation in the extracellular near-membrane concentration traces. For an optional exponential fit,

$$C_i(t) = C_{i,\infty} - \Delta C_i e^{-t/\tau_i}, \tag{S15}$$

where $C_{i,\infty}$, $\Delta C_i$, and $\tau_i$ are fitted parameters. The code accepts a fit when $R^2 > 0.98$, $\tau_i > 0$, and $\tau_i < 5T_{\mathrm{sim}}$. Otherwise, the fitted timescale is reported as bounded by the simulated interval.

For model-free analysis, fractional relaxation is defined relative to the observed change within the sampled window:

$$u_i(t) = \frac{C_i(t) - C_i(0)}{C_i(T_{\mathrm{sim}}) - C_i(0)}. \tag{S16}$$

Linear interpolation locates the first crossings of $u_i = 0.1$, 0.5, and 0.9 and thereby determines $t_{10}$, $t_{50}$, and $t_{90}$. The integrated measure is

$$\tau_{\mathrm{int},i} = \int_0^{T_{\mathrm{sim}}} [1 - u_i(t)]\, dt. \tag{S17}$$

These measures are defined relative to the sampled endpoint, rather than a demonstrated equilibrium concentration. In particular, $\tau_{\mathrm{int},i}$ should not be interpreted as a converged relaxation eigenmode timescale when the trajectory has not reached its long-time state.

#### 3. Three-dimensional pore-resolved model

##### 3.1 Computational domain and pore geometry

The three-dimensional model explicitly resolves the pore geometry within a finite Cartesian domain:

$$[-L_x/2, L_x/2] \times [-L_y/2, L_y/2] \times [-L_c, L_w].$$

The default values are $L_x = L_y = L_c = L_w = 1000$ nm. The nominal membrane thickness is $L_M = 4$ nm. Membrane grid planes satisfy $|z| \le L_M/2 + \Delta z_{\mathrm{min}}/2$, accommodating the finite mesh spacing.

A cylindrical aqueous region spanning the membrane represents each pore. The default pore radius is $R^P = 10.75$ nm. Membrane cells outside the pores have lipid properties, whereas pore cells retain aqueous dielectric properties.

The grid is stretched geometrically away from the origin. Default minimum spacings are $\Delta x_{\mathrm{min}} = \Delta y_{\mathrm{min}} = 0.5$ nm and $\Delta z_{\mathrm{min}} = 0.2$ nm. Growth ratios are 1.12 laterally and 1.10 axially, and maximum spacings are 10 nm laterally and 40 nm axially. Adjacent face spacings determine the finite-volume widths.

The accessibility mask excludes protein centers from the lipid membrane and its radius-dependent axial margin. Within a cylindrical pore, the nominal accessible radius is

$$R_{\mathrm{access},p} = \max\left(R^P - a_p, 0\right). \qquad \text{(S18)}$$

Protein-specific transport masks enforce this accessibility condition. Proteins with radii approaching the pore radius have progressively restricted accessible regions. No Henry-function pore-hindrance factor is applied in the 3D transport solver.

##### 3.2 Protein diffusivity and concentration-dependent crowding

The default protein species has $C_p = 1$ mM, $z_p = -10$, and $a_p = 5$ nm. Multiple protein species with independently configured concentrations, radii, and valences are supported.

The model calculates an aqueous Stokes–Einstein diffusivity and applies an additional threefold reduction:

$$D_{p,c} = \frac{1}{3}\frac{k_B T}{6\pi\eta_w a_p}, \qquad \eta_w = 0.6913\ \mathrm{mPa\,s}. \qquad \text{(S19)}$$

For the default protein, $D_{p,c} \approx 2.19 \times 10^{-11}$ m² s⁻¹. The factor of one-third is a prescribed phenomenological reduction of the aqueous diffusivity. Although denoted $D_{p,c}$ in the code, this baseline coefficient is assigned throughout protein-accessible aqueous regions. It is subsequently modified by crowding and geometric accessibility during transport.

A spatial convolution of protein number density gives the local occupied-volume fraction:

$$\varphi(\mathbf{r}) = \sum_p V_p \left(K_p * n_p\right)(\mathbf{r}), \qquad V_p = \frac{4\pi a_p^3}{3}, \qquad \text{(S20)}$$

where $K_p$ is a normalized spherical averaging kernel of radius $a_p$, and $V_p$ is the corresponding particle volume. For the default species, the maximum permitted occupied fraction is $\varphi_{\max} = 0.64$.

The concentration-dependent transport gate $g(\varphi)$, defined in Eq. (7), multiplies both ionic and protein effective diffusivities in the 3D time-advancement kernel. The maximum permitted packing concentration also limits protein number density.

Ion and protein transport follow the Nernst–Planck structure in Eqs. (2) and (8), respectively, with the corresponding crowding and geometric-accessibility factors. Because the same effective diffusivity enters diffusion and electromigration, the 3D transport law satisfies the conventional Nernst–Einstein relation. No separate Henry-corrected electrophoretic mobility or additional permittivity multiplier is used.

The 3D charge density is

$$\rho_{\mathrm{total}} = F\left(\sum_i z_i C_i + \sum_p z_p \frac{n_p}{N_A}\right) + \rho_{\mathrm{fixed}} + \rho_{\mathrm{wall}}, \qquad \text{(S21)}$$

where $\rho_{\mathrm{fixed}}$ and $\rho_{\mathrm{wall}}$ denote the optional immobile volume charge and pore-wall charge contributions. The physiological ions and charged auxiliary species X contribute to the ionic sum; neutral species Y contributes only to osmolarity.

##### 3.3 Initial conditions and auxiliary solutes

The model initializes concentrations using a smooth axial interpolation between the cytosolic and extracellular values:

$$a(z) = \frac{1}{2}\left[1 - \tanh\left(\frac{z}{2L_M}\right)\right]. \quad \text{(S22)}$$

For each physiological ion,

$$C_i(\mathbf{r}, 0) = \left[a(z)C_{i,c} + \left(1 - a(z)\right)C_{i,w}\right]m_i(\mathbf{r}), \quad \text{(S23)}$$

where $m_i$ is the aqueous accessibility mask. Protein is initialized analogously using its species-specific accessibility mask and is stored as number density.

At each grid point, the model initializes charged auxiliary species X to cancel the net local charge from physiological ions, protein, and any configured immobile charge, subject to nonnegative concentrations. It then adds neutral species Y to make the total initial particle concentration approximately uniform across accessible voxels. The simulation stores both auxiliary fields alongside the physiological ions and evolves them as mobile species.

In the 3D implementation, X and Y each have diffusivity $D_{XY} = 1.0 \times 10^{-9}$ m² s⁻¹. An optional fixed charged species can be configured separately and contributes to both the charge density and osmolarity. The default configuration disables this species.

##### 3.4 Membrane potential and pore-wall charge

The model imposes the resting potential by transferring physiological ions between thin aqueous slabs adjacent to the membrane. Transfer weights scale with $z_i^2 \overline{C}_i$, and the procedure excludes X and Y.

The slab thickness is $\max(\lambda_D, 1.5\Delta z_{\min})$, where $\lambda_D$ is calculated from the ionic concentrations and protein charge contribution. The target potential is −75 mV. Bracketing and bisection determine the shift magnitude from the Poisson-resolved difference between volume-weighted slab potentials. The default acceptance tolerance is 0.05 mV.

The model represents pore-wall charge using a shell around the nominal cylindrical pore surface. For $N_P$ pores with radius $R^P$ and membrane thickness $L_M$, the configured surface charge corresponds to total wall charge

$$Q_{\text{wall}} = \sigma N_P \, 2\pi R^P L_M. \quad \text{(S24)}$$

When present, the wall-charge field distributes this total charge over the discrete wall-shell volume. A spherical weighting kernel can introduce oppositely charged counterions within a local screening region. The Debye length and minimum mesh spacings determine the kernel radius.

During construction of static-pore cases A–C, the supplied implementation sets the initial wall-charge array to zero and does not invoke the surface-charge reconstruction function for those geometries. Consequently, a nonzero configured $\sigma$ does not by itself establish a nonzero wall-charge field in those cases. Dynamic pore insertion constructs the wall-charge field when it rebuilds the geometry. This distinction is relevant when interpreting surface-charge-dependent simulations.

##### 3.5 Numerical discretization and Poisson solver

The 3D model uses a finite-volume formulation on the nonuniform Cartesian grid. Harmonic means define face permittivities and diffusivities, while a three-dimensional Scharfetter–Gummel scheme discretizes species fluxes.

Discretizing Poisson's equation produces a symmetric positive-definite linear system. A preconditioned conjugate-gradient algorithm solves this system, and its preconditioner combines batched tridiagonal solves along the three Cartesian directions. The axial outer boundaries use $\phi = 0$, while lateral potential boundaries use zero normal gradient. The default relative residual tolerance is $10^{-5}$, with a floor of $3 \times 10^{-6}$ in single precision. Extrapolation from preceding potential fields initializes subsequent Poisson solves.

Explicit Euler steps advance transport. A fourth-order series evaluates the Scharfetter–Gummel Bernoulli function for small arguments, while asymptotic forms handle large arguments. Missing neighbors at the outer boundaries contribute zero flux.

An explicit diffusion stability estimate is

$$\Delta t_{\mathrm{diff}} = \frac{0.45}{2D_{\max}\left(\Delta x_{\min}^{-2} + \Delta y_{\min}^{-2} + \Delta z_{\min}^{-2}\right)}. \quad \text{(S25)}$$

The stability cap limits the maximum step to $0.9\Delta t_{\mathrm{diff}}$. The simulation holds the default initial step of 2.5 ps fixed through the configured resolution interval. Thereafter, the step increases geometrically by a factor of 1.03 without exceeding the cap. When necessary, shorter steps align the integration with output times or pore-formation events.

CuPy implements arrays, convolution operations, and custom transport kernels on an NVIDIA GPU. Field arrays use float32 by default, whereas selected inner products, integrated fluxes, and reductions use float64. Direct evaluation handles small protein-density kernels, and FFTs handle larger kernels.

##### 3.6 Pore configurations and sequential opening

The pore geometries comprise four principal cases. Case A contains a single central pore. Case B arranges the specified number of pores on an approximately square centered grid. Case C shifts the grid to one side of the computational domain. Case D begins with a central seed pore and subsequently inserts additional pores at nonoverlapping positions within a configured spatial region.

The simulation inserts pores either at predetermined times or in response to the potassium-efflux feedback condition. Let $J_K(t)$ denote the total $K^+$ molar flow across the membrane diagnostic plane, with positive sign corresponding to outward transport. Cumulative positive potassium efflux is

$$Q_K^+(t) = \int_0^t \max\,[J_K(t'), 0]\, dt'. \quad \text{(S26)}$$

For a prescribed threshold $Q_K^*$, the simulation inserts additional pores when

$$Q_K^+(t) \geq mQ_K^*, \qquad m = 1,2, \ldots, \quad \text{(S27)}$$

until the configured number of pores has been reached.

After pore insertion, the code updates the pore, lipid, wall, permittivity, and species-accessibility fields. Prescribed axial concentration profiles initialize newly opened cells, subject to protein accessibility and packing constraints. The update then adjusts local chloride concentration to restore electroneutrality and reconstructs the Poisson operator for the new geometry.

##### 3.7 Flux, electrical, and osmotic diagnostics

The model evaluates transmembrane fluxes at the discrete axial face nearest $z = 0$. Positive flux denotes transport from the cytosolic to the extracellular side. The following expression gives the total electrical current:

$$I = F\left[\sum_i z_i \int_{z=0} \mathbf{j}_i \cdot \hat{\mathbf{z}}\, dA + \sum_p z_p \int_{z=0} \mathbf{j}_p^{(\mathrm{mol})} \cdot \hat{\mathbf{z}}\, dA\right], \qquad \text{(S28)}$$

where $\mathbf{j}_p^{(\mathrm{mol})} = \mathbf{j}_p / N_A$ is the protein molar flux corresponding to the number-density formulation.

The difference between volume-weighted potential averages in aqueous slabs adjacent to the membrane defines the membrane potential. For local intracellular and extracellular osmolarities, the initial axial osmolarity gradient sets the sampling distance from the membrane.

The code records ionic and protein fluxes, cumulative positive efflux, protein leakage, total charge, pore number, Poisson solver iterations, and selected three-dimensional fields. An optional quiescence diagnostic assesses changes in cumulative efflux over repeated output intervals after prescribed minimum time and transport thresholds. Satisfying this criterion generates a notification but does not terminate the simulation.

In the supplied code, the ion-flux diagnostic at the membrane plane does not include the same concentration-dependent crowding multiplier used by the ion time-advancement kernel. Consequently, diagnostic flux values, including those used to accumulate potassium efflux, are not in general identical to the crowding-modified fluxes used for concentration updates. This implementation difference must be considered when interpreting quantitative flux and feedback measurements.

#### 4. Code availability

The Python implementations are available in the 1D_model/ and 3D_model/ directories of the accompanying repository:

https://github.com/c1462881/Multispecies-transport-model

The 1D implementation uses Python, NumPy, SciPy, pandas, and Matplotlib and is configured through configuration.py. The original model is executed using python main.py.

The 3D implementation additionally requires CuPy, a compatible CUDA installation, and an NVIDIA GPU. YAML files specify simulation-specific parameters, and python main.py --config config.yaml launches the model. The YAML configuration and output records are required to establish the exact parameters used for individual manuscript figures.

#### Figures

**Classical Donnan versus multispecies transport scenarios**

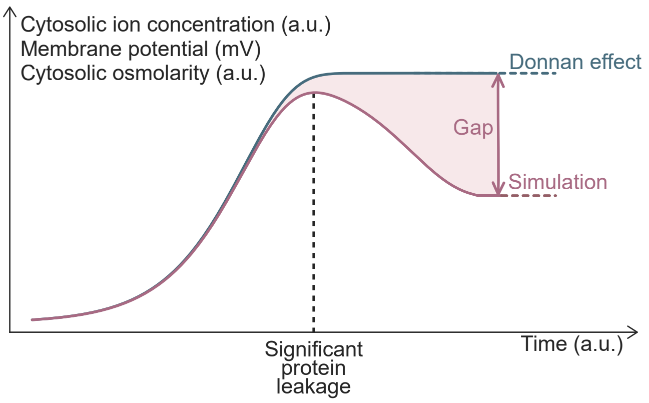

**Figure S1. Protein transport enables deviation from Donnan equilibrium.**

**A Extended temporal evolution of concentrations, electric potential, and osmolarity**

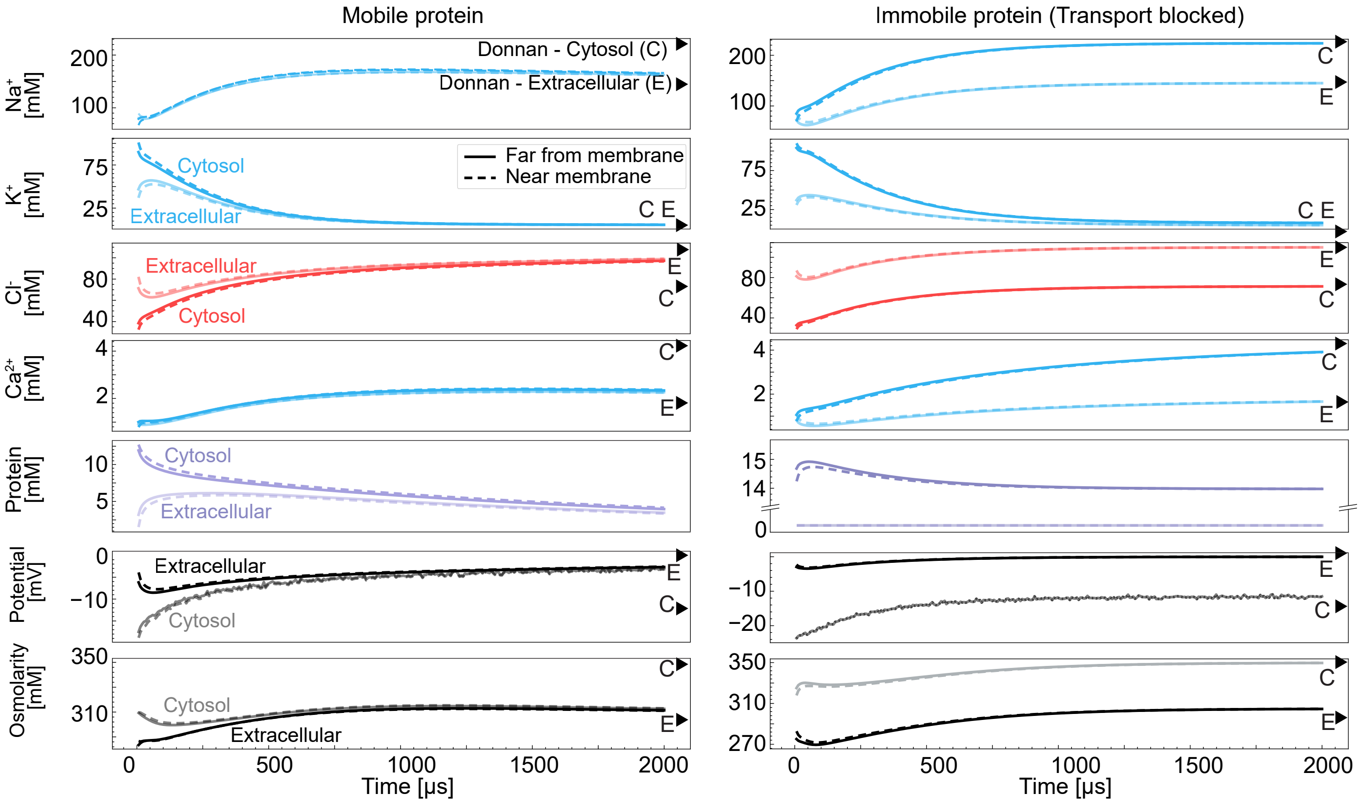


**B Final spatial profiles at extended time for mobile versus blocked protein**

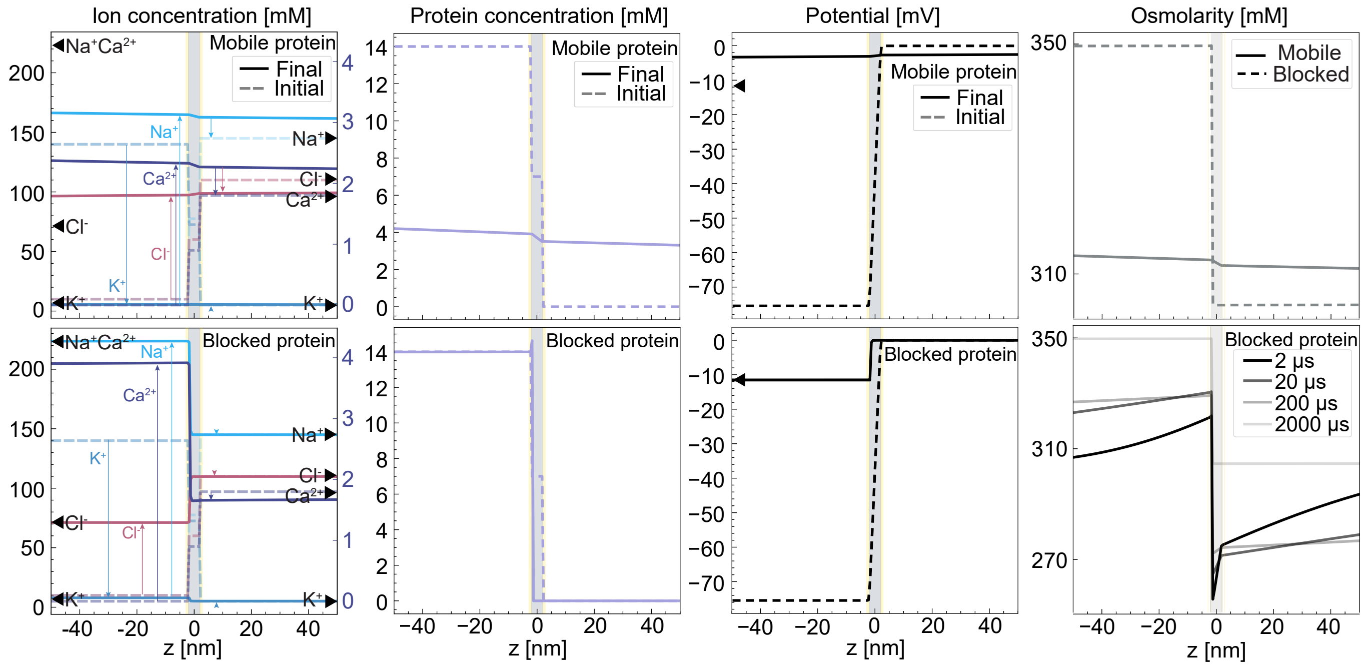

**Figure S2. Multispecies transport distinguishes from Donnan over extended time.**

(A) Mobile proteins progressively deviate from the Donnan state, while blocked proteins sustain the full Donnan ion distribution, potential offset, and osmotic imbalance.

(B) A mobile-protein system eventually reaches a balanced distribution of all species, while a blocked-protein system eventually settles onto the exact Donnan state.

**Time-resolved electric potential maps of a single pore simulation**

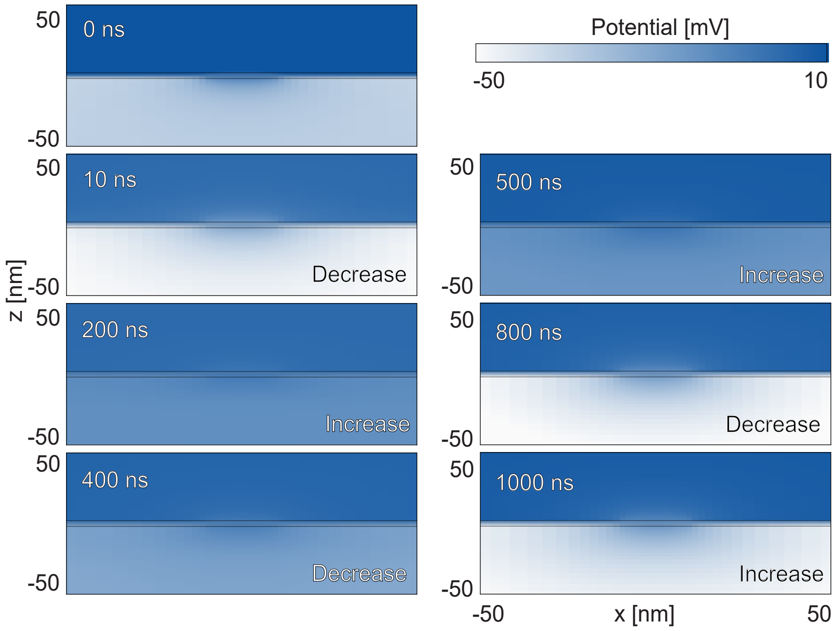

**Figure S3. Electric potential evolves dynamically around a pore.**

**Osmolarity trajectories at fixed distances from the membrane**

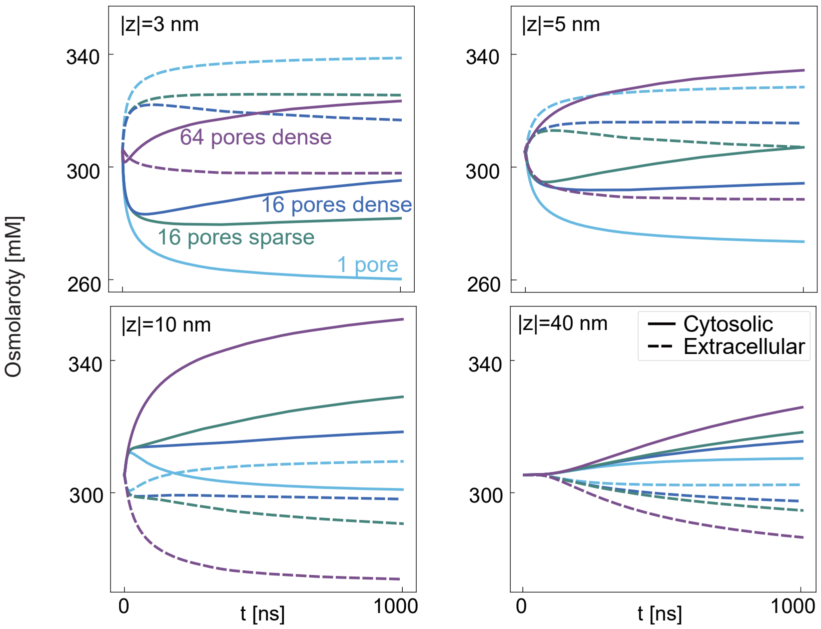

**Figure S4. Position-resolved osmolarity varies across pore clustering configurations.**

**Spatiotemporal evolution with immobile background species**

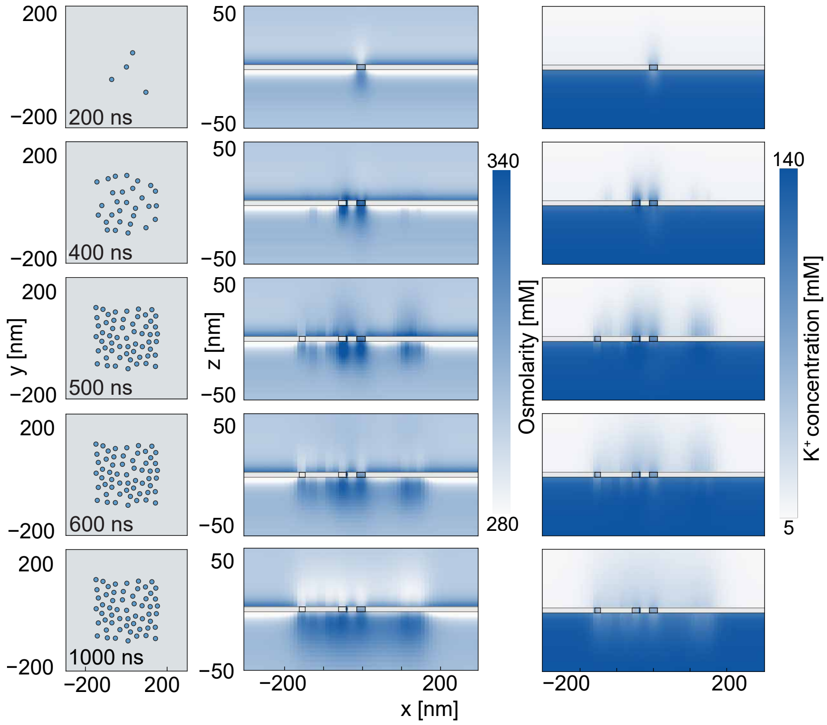

**Figure S5. Immobility of background species strengthens positive feedback effects.**
